# deepBlink: Threshold-independent detection and localization of diffraction-limited spots

**DOI:** 10.1101/2020.12.14.422631

**Authors:** Bastian Th. Eichenberger, YinXiu Zhan, Markus Rempfler, Luca Giorgetti, Jeffrey A. Chao

## Abstract

Detection of diffraction-limited spots is traditionally performed with mathematical operators designed for idealized spots. This process requires manual tuning of parameters that is time-consuming and not always reliable. We have developed deepBlink, a neural network-based method to detect and localize spots automatically and demonstrate that deepBlink outperforms state-of-the-art methods across six publicly available datasets. deepBlink is open-sourced on PyPI and GitHub (https://github.com/BBQuercus/deepBlink) as a ready-to-use command-line interface.

---

In biomedical research, the detection, counting, and localization of sub-diffraction fluorescent signals (spots) represent essential steps in various imaging applications including particle tracking in live cell imaging data and quantification of mRNAs in single-molecule fluorescent *in situ* hybridization (smFISH). While advances in smFISH-based spatial transcriptomics have enabled the quantification of thousands of mRNAs in single cells, these methodologies have focused on the development of transcript barcoding strategies for multiplexing and have relied on conventional threshold-based detection of single mRNA molecules [1, 2]. Accurate, high-throughput spot detection and localization in single-molecule fluorescent microscopy images with varying background brightness levels and spot qualities, however, poses a challenge for current spot detection methods. Current methods such as the broadly adopted TrackMate [3] are based on intensity thresholds and rely on mathematical operators (e.g. Laplacian of Gaussian) that require ad-hoc adjustments of parameters on a cell-by-cell and image-by-image basis. Fully automated, user-friendly, and accurate spot detection and localization methods are currently not available.

Recent advances in deep learning have consolidated the convolutional neural network (CNN) as the state-of-the-art for computer vision applications [4, 5]. Previously, CNNs have been used for threshold-independent particle detection [6, 7, 8], however, without localizing particle centers with sub-pixel resolution. This prevents accurate positional measurements and thus their application in high-resolution microscopy. In addition, while the source-code of some methods is available, none of them have been made easily accessible for non-experts.

Here we present deepBlink, a simple command-line interface that exploits neural networks to detect and localize diffraction-limited spots in microscopy images. deepBlink can batch process images with different background brightness levels and spot qualities without relying on manual adjustments of thresholds. deepBlink’s interface provides the ability to detect and localize spots on images using a pre-trained model (Figure 1 Steps 5-9). Pre-trained models are available for download using deepBlink (Figure 1 Step 4). Alternatively, new models can also be trained in deepBlink using custom-labeled data (i.e. images and their corresponding spot-coordinate annotations) (Figure 1 Steps 1-3). To customize training, the plug-and-play software architecture with a central configuration file allows more experienced deep learning practitioners to easily adjust existing functionalities or add new ones. Instructions on how to install and use deepBlink are described in a tutorial (https://youtu.be/vlXMg4k79LQ).

**Figure 1:**
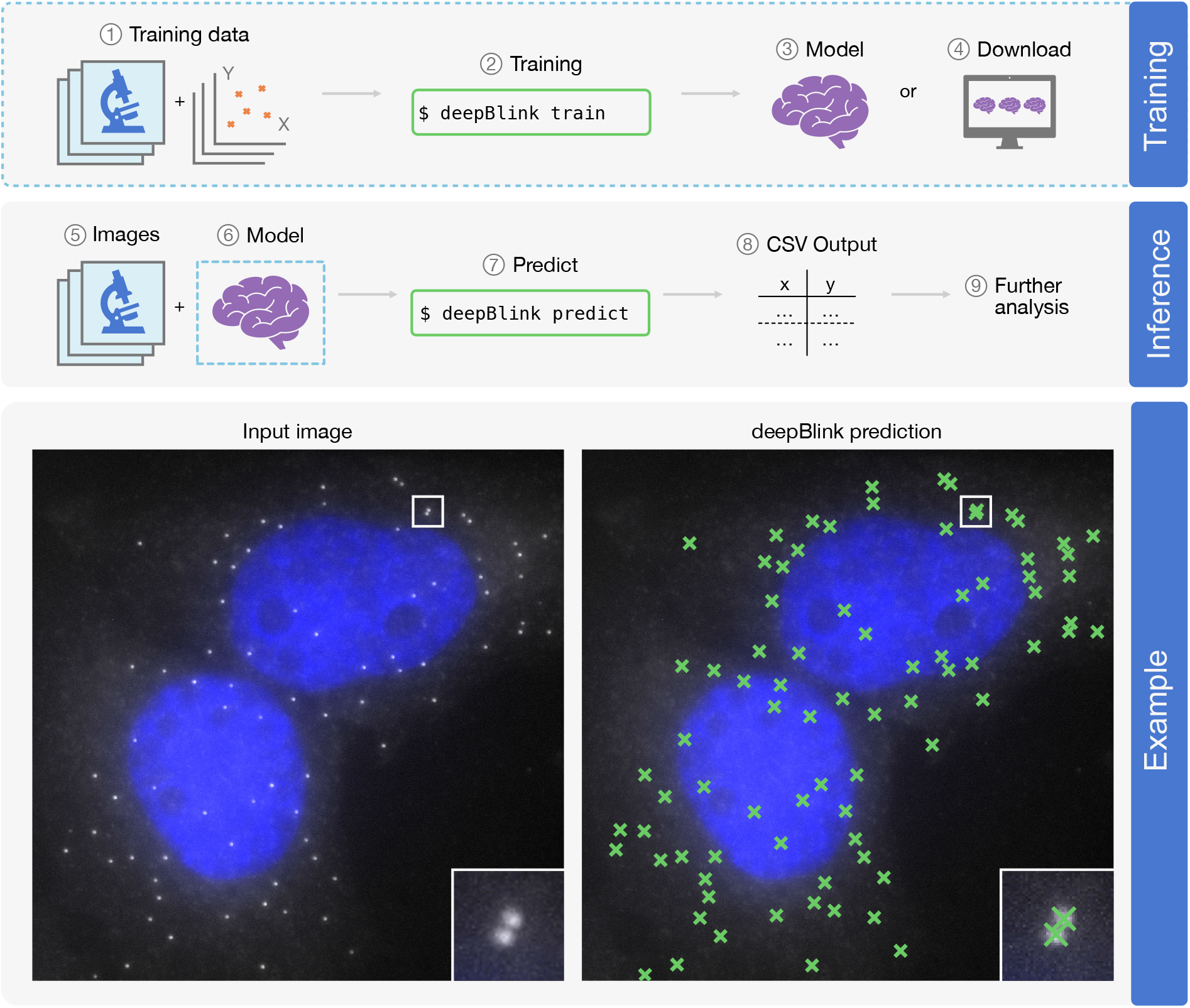
Overview of deepBlink’s functionality. deepBlink requires a pre-trained model that can be obtained by training from scratch using custom images and coordinate labels (1-3) or downloaded directly (4). To predict on new data, deepBlink takes in raw microscopy images (5) and the aforementioned pre-trained model (6) to predict (7) spot coordinates. The output is saved as a CSV file (8) which can easily be used in further analysis workflows (9). An example use case is shown for a smFISH analysis with blue indicating DAPI staining.

The current default network architecture we propose is a fully convolutional neural network that exploits the U-Net architecture [9] to encode features that are used for spot coordinate prediction (Supplementary Figure 1). The network takes in microscopy data and maps the input images onto smaller regions we termed “grid-cells”. For each grid-cell, the network predicts the probability and localization of a spot within its region. This process works similarly to the region-based bounding-box predictions originally developed in the YOLO architecture [10].

Each grid-cell returns three values corresponding to the probability, the x, and the y position of a single spot. The performance of the network critically depends on the cell-size (width/height in pixels of one grid-cell in the original image) as too small sizes can lead to a class imbalance. Given a fixed number of spots in an image, decreasing the cell-size results in fewer grid-cells containing a spot. This will create an imbalance for the classification part of the loss, favoring background (grid-cells without spots). Images with higher spot densities perform better even at smaller cell-sizes (Supplementary Figure 2). In contrast, bigger cell-sizes increase the risk of one grid-cell containing multiple spots that pose the known design limitation as one grid-cell cannot regress to multiple coordinates [11]. We found a cell-size of four to be ideal for most scenarios, as only around 0.0003% of grid-cells contained more than one spot across all datasets used (Supplementary Figure 2). Although a cell-size of four was found to be optimal, datasets containing significantly more or less dense regions of spots could require smaller or larger cell-sizes respectively. deepBlink allows for the configuration of cell-sizes when training new models.

The basic building blocks of our proposed architecture consist of three convolutional layers followed by a squeeze and excitation layer [12]. We employ spatial dropout layers [13] before every downsampling step to increase training stability and to reduce overfitting. We trained the models by minimizing the dice loss for the classification (J_class_, Equation 1) and the average root mean square error (RMSE) across all spots (n) for the localization (J_loc_, Equation 2). The objective function (J) is shown in Equation 3:

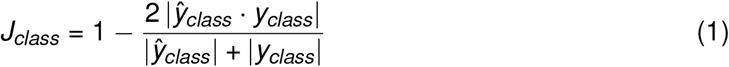

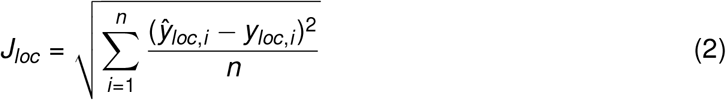

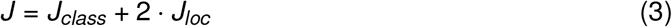

Ablation experiments (Supplementary Table 2) showed that the most important factors for model performance were the usage of dice loss and the addition of squeeze blocks. We experimented with other architectural blocks, namely inception [14] and residuals [15], that have been employed in similar computer vision tasks [16] but were not able to observe any significant improvements. We found that having a constant number of filters (64) across every layer of the network performs better than the typical increase/decrease in filters for every downsampling/upsampling step, respectively. This suggests that higher-level features found in deeper layers are less important for spot detection.

To compare the performance of deepBlink to previously published methods, we used a single metric we termed the “F1 integral score” (see Supplementary Figure 3 and Methods) that takes into account both spot detection and spot localization. The F1 integral score ranges between zero and one where one corresponds to perfect prediction and zero denotes that no prediction is within a 3px radius around the ground truth spot positions.

To measure performance on a variety of spot sizes, shapes, densities, and signal-to-noise ratios (SNRs), we used two real, manually labeled microscopy datasets (smFISH [17] and Sun-Tag live cell single particle tracking [18]) and four synthetic datasets (a custom Particle dataset generated using [19] and the Microtubule, Vesicle, and Receptor datasets from the International Symposium on Biomedical Imaging (ISBI) particle tracking challenge [20]). Exemplary images of each dataset are displayed in Supplementary Figure 4. Since manual labeling is time-consuming, we measured how many images have to be labeled until only marginal performance increases are observed by adding more images. Based on the F1 integral score, a training set size of around 100 images is enough to achieve sufficient prediction quality on datasets with narrow SNR distributions (Supplementary Figure 5a,b), while more images are required for datasets with broader distributions (Supplementary Figure 5a,c).

On average, deepBlink is able to outperform other methods on all datasets (Table 1), both in detection (average efficiency above 85%, Supplementary Table 3) and in localization (average error below 0.5 pixels, Supplementary Table 4). Specifically, we compared the performance of deepBlink against a popular classical method and a state-of-the-art deep learning-based method. TrackMate [3] is a Fiji plugin that uses the Laplacian of Gaussian (LoG) to detect and localize spots. This requires the setting of manual thresholds. DetNet [8] is a more recent deep learning-based method which has been shown to perform well on the synthetic ISBI datasets. To improve the fairness of the comparison, we optimized hyperparameters for both benchmarking methods (TrackMate and DetNet) on every dataset individually (see Methods). While these methods can detect most spots, deepBlink still outperforms other softwares without dataset-specific tuning, which will make deepBlink more accessible for a large variety of applications. Additionally, we fitted a Gaussian function on the original input data initialized by deepBlink’s prediction output and did not observe any improvement in localization (Supplementary Table 4). This further demonstrates deepBlink’s localization precision.

**Table 1:** F1 integral score (mean ± standard deviation) results for three methods across six datasets. Statistical significance was determined by one-sided Wilcoxon signed-rank test (deepBlink greater than) with **: p < 1e^−2^, ****: p < 1e^−4^. Size of test set: 129 smFISH, 105 SunTag, 64 Particle, 240 Microtubule, 240 Receptor, and 240 Vesicle.

|  |  | TrackMate |  | DetNet |  | deepBlink |
| --- | --- | --- | --- | --- | --- | --- |
| Real | smFISH | 0.865 ± 0.177 | **** | 0.442 ± 0.250 | **** | **0.905 ± 0.145** |
| | SunTag | 0.652 ± 0.328 | ** | 0.036 ± 0.040 | **** | **0.712 ± 0.279** |
| Synthetic | Particle | 0.941 ± 0.008 | **** | 0.828 ± 0.014 | **** | **0.944 ± 0.008** |
| | Microtubule | 0.380 ± 0.228 | **** | 0.298 ± 0.149 | **** | **0.637 ± 0.261** |
| | Receptor | 0.606 ± 0.344 | **** | 0.517 ± 0.258 | **** | **0.682 ± 0.310** |
| | Vesicle | 0.459 ± 0.198 | **** | 0.520 ± 0.245 | **** | **0.732 ± 0.259** |
| Average | | 0.651 ± 0.214 | | 0.440 ± 0.159 | | **0.769 ± 0.210** |

To determine the effect of spot densities and SNRs on detection performance, we used the official categories from the ISBI particle tracking challenge and divided the Microtubule, Receptor, and Vesicle datasets accordingly (see Methods). In general, both spot density and SNR does not affect the performance of deepBlink except for very low SNRs where no model is able to perform well (Supplementary Figure 6, Supplementary Figure 7, and Supplementary Table 5).

In summary, deepBlink enables automated detection and accurate localization of diffraction-limited spots in a threshold-independent manner. deepBlink is packaged in an easy and ready to use command-line interface, allowing both novel and expert users to efficiently analyze their microscopy data. All pre-trained models are available for download from Figshare (see Methods) or through the command-line interface. We benchmarked deepBlink on six publicly available datasets that contain both synthetic images as well as smFISH and live cell (SunTag) imaging data, each with a variety of spot sizes, shapes, densities, and signal-to-noise ratios. We show that deepBlink outperforms all benchmark methods, both in detection and localization across these datasets. We expect that our improved detection efficiency and localization precision will also enhance current particle tracking analyses and make image processing more reliable and reproducible.

## Methods

### Training data

Datasets “Microtubule”, “Receptor”, and “Vesicle” were created from the ISBI Particle Tracking Challenge 2012 [20]. Each dataset consisted of 1200 images and was split into training/-validation/test splits ensuring equal distribution of signal to noise ratios and densities. The synthetic “Particle” dataset was generated using the “Synthetic Data Generator” Fiji plugin (smal.ws/wp/software/synthetic-data-generator/) described in Smal et al. [19]. A pool of 576 images was created with various signal to noise ratios and background intensities to mimic small diffraction-limited particles. The “smFISH” dataset was created from a pool of 643 manually annotated smFISH images originally published in Horvathova et al. [17]. The “SunTag” dataset consists of 544 manually annotated live cell SunTag images originally published in Mateju et al. [18].

### Dataset labeling

Labeling for both real datasets was performed using TrackMate. Each image was subjected to manual thresholding and falsely detected spots were manually corrected. Labels were exported using the “All Spots statistics” tool. Once all images were labeled, the deepBlink command deepBlink create was used to convert all images and labels into a partitioned dataset file.

### Network architecture

deepBlink’s default network is based on the U-Net architecture [9]. We used a second encoder to reduce the fully sized representation into a smaller one corresponding to the cell-size. This second encoder is variable in its depth to adjust for the configured cell-size. Each encoding step consists of a convolutional block, squeeze and excitation block, and spatial dropout (Supplementary Figure 1 and Supplementary Table 1). Decoding steps had the same layout but did not employ dropout. The architecture was implemented in Python version 3.7 using the TensorFlow framework version 2.2.

### Network training

All deepBlink trainings were performed using the public command-line interface on version 0.1.0 on the default settings provided by deepBlink config. In particular, we used a cell-size of 4, a dropout rate of 0.3, a batch size of 2, a learning rate of 0.0001, and the AMSGrad stochastic optimizer [21]. Training was run on a computer cluster with the following specifications: 64-bit CentOS Linux 7 (Core), 96 cores (3926 threads) Intel(R) Xeon(R) Platinum 8168 CPU 2.70GHz, 1007 GB RAM, 8 Nvidia GeForce RTX 2080 Ti GPUs (11 GB VRAM). Training on average took around 5 hours per model. All models were trained for 200 epochs.

### Benchmarking of TrackMate

TrackMate [3] is not a machine learning method and relies on manual thresholding. To find the best parameters, we tuned TrackMate on the pool of training and validation images from each dataset. More specifically, we performed a grid search across different values of “Estimated blob diameters” in the “LoG” filter option in Fiji. Subsequently, the resulting quality scores were normalized to the median of all images. Twenty linearly distributed quantiles were applied. At each quantile, the F1 integral score was calculated for each image. The finally selected parameters (blob diameter and absolute quantile value) maximized the mean F1 score on the combined training and validation dataset.

### Benchmarking of DetNet

Due to the lack of a publicly available implementation of DetNet [8], we implemented the model and followed the training procedure in the publication. Notably, the number of parameters in our model was 10x larger than originally described. Using the coordinate locations, ground truth output masks were created with a single pixel of value one at the centroid position. The output segmentation maps were converted back to coordinates by binarizing and taking the centroids of each fully connected component. A hyperparameter search was performed across twenty linearly distributed sigmoid shifts *α*. The *α* with the highest validation F1 score was used for evaluation on the test set. Supplementary Table 6 lists the *α* values used for each dataset.

### Benchmarking of deepBlink

deepBlink was trained through the command-line interface version 0.1.0. All datasets were processed individually but without dataset-specific hyperparameter optimization. The models were then evaluated on the corresponding test sets.

### F1 integral score

The F1 integral score was selected as a metric to compare the detection and localization accuracy of given models. A linear distribution of 50 cutoff values was created between 0px and 3px. At each cutoff value, ground truth and predicted coordinates with Euclidean distances below the given cutoff were matched using the Hungarian method [22]. Finally, the trapezoidal rule was used to integrate the F1 score curve (Supplementary Figure 3). The integrated value was then normalized to the maximum area resulting in a minimum score of 0 and a maximal score of 1. An algorithmic overview is described in Algorithm 1 of which an implementation can be found in `deepBlink’s source code under deepblink.metrics.f1_integral`.

**Algorithm 1:**
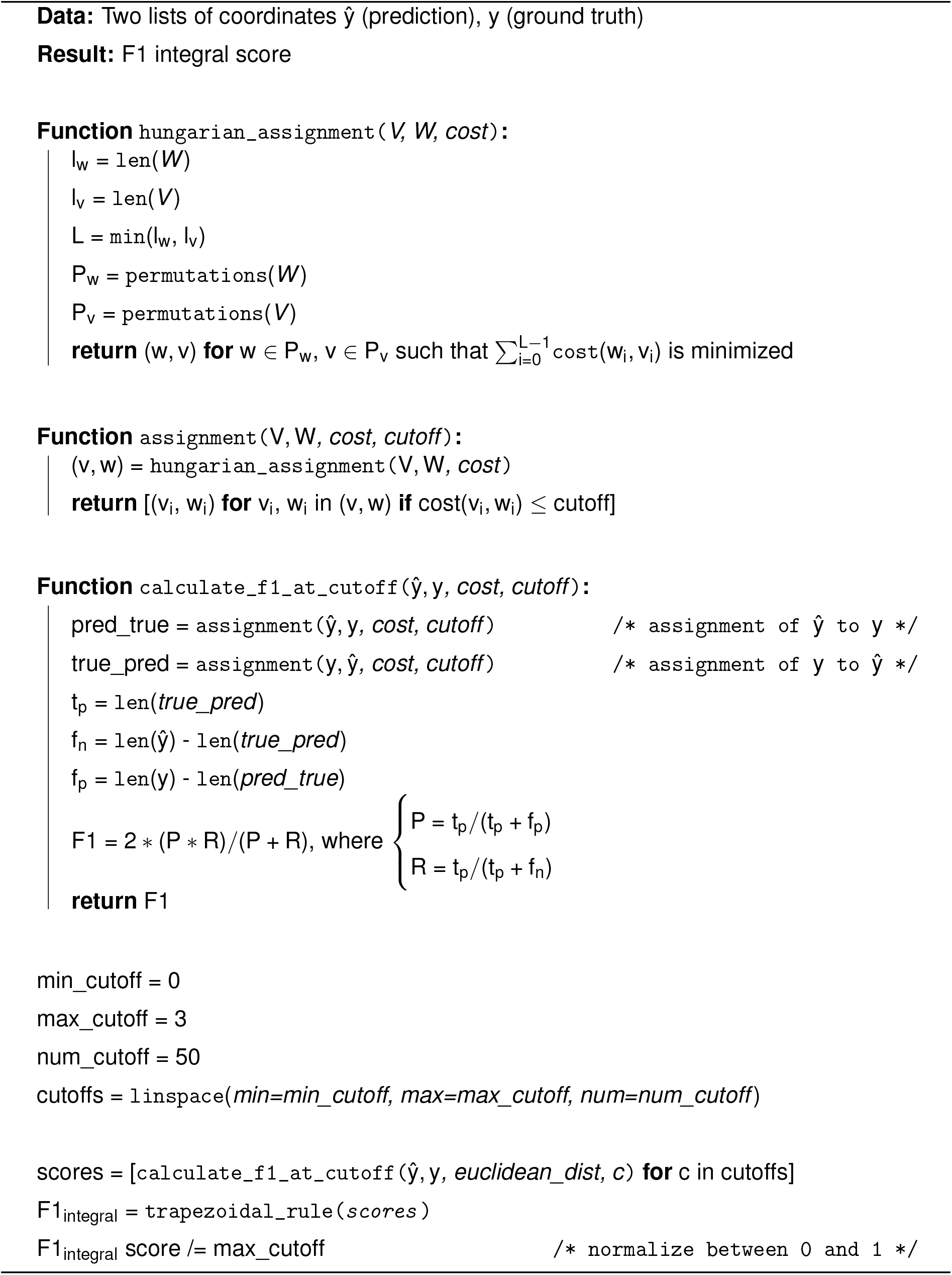
Algorithmic description of the F1 integral score

### Root mean square error

The root mean square error (RMSE) was defined as the mean Euclidean distance of all true positive coordinates at a cutoff value of 3px. True positive coordinates are predicted coordinates with an assignment to a ground truth coordinate.

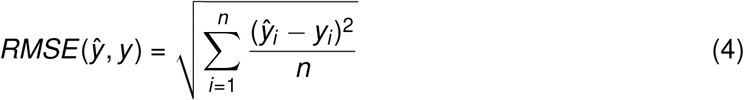

## Data availability

All datasets with their training, validation, and test splits are available on Figshare (https://figshare.com/projects/deepBlink/88118) together with all trained models used for bench-marking. Further data will be made available from the corresponding authors upon request.

## Code availability

deepBlink’s command-line interface is available on PyPI. Updated versions of the software can also be found on GitHub https://github.com/BBQuercus/deepBlink.

## Acknowledgements

The authors thank Guido Tiana for great ideas in training optimization, Jan Eglinger for support in benchmarking for TrackMate, and Gregory Roth for mathematical advice. We acknowledge the Computational Biology platform at the FMI for providing the computational resources required. We thank Ivana Horvathova / Franka Voigt and Daniel Mateju for providing the smFISH and SunTag images respectively. We thank members of the Giorgetti and Chao laboratories for assistance in preliminary data labeling. Research in the Giorgetti laboratory is funded by the Novartis Foundation and the European Research Council (ERC) under the European Union’s Horizon 2020 research and innovation (grant agreement no. 759366, “BioMeTre”). Research in the Chao laboratory is funded by the Novartis Foundation, the Swiss National Science Foundation (grant no. 31003A_182314), and the SNF-NCCR RNA & Disease.

## Author contributions

B.T.E. and Y.Z. coded the software and trained models. B.T.E. coded the benchmarking routines and conceived the idea. B.T.E and Y.Z. directed the project and wrote the paper. M.R., L.G., and J.A.C jointly supervised this work. All authors read the paper and approved the content.

## Supplementary Information

**Supplementary Figure 1:**
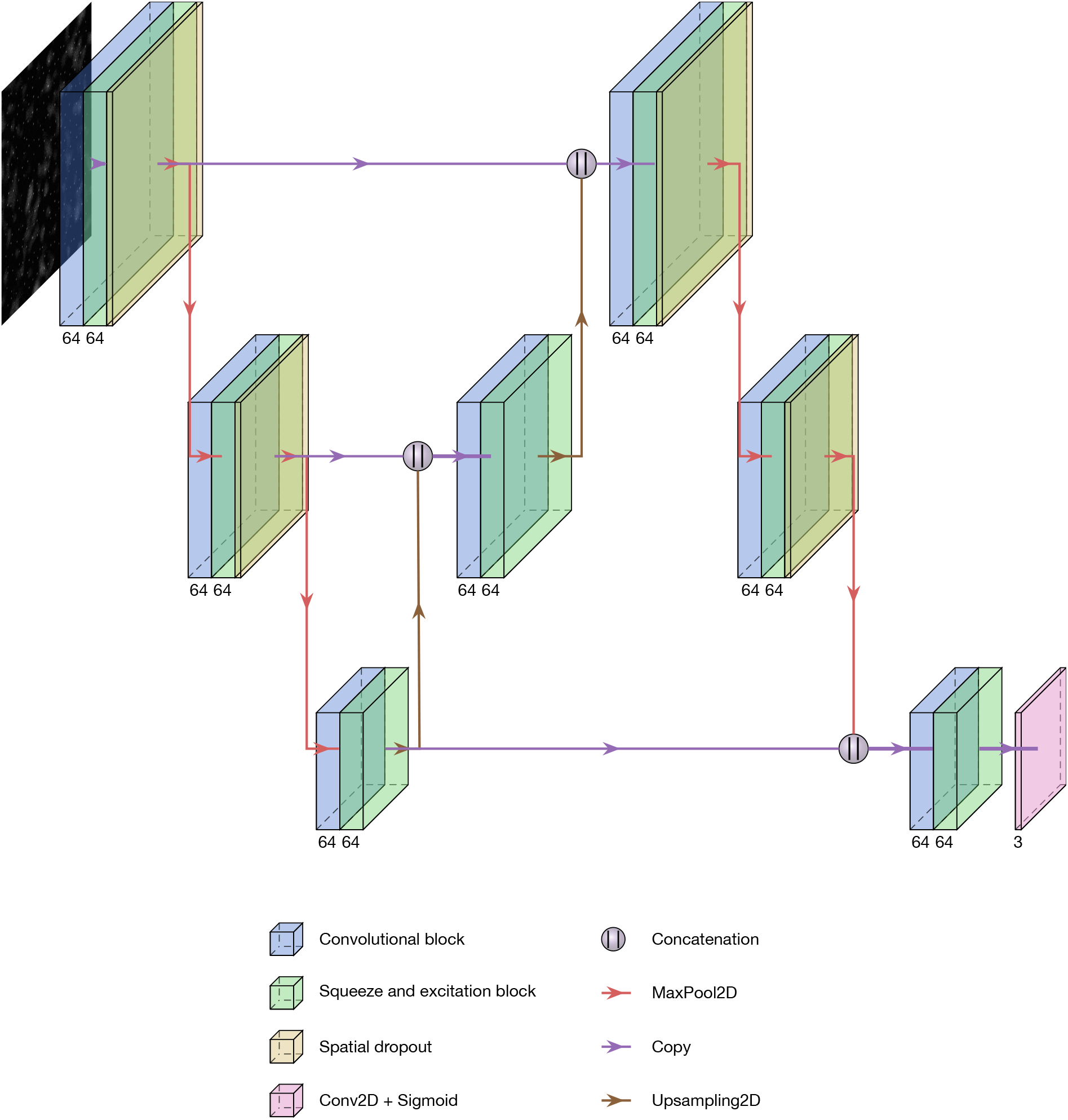
Overview of the default neural architecture for deepBlink (components are described in Supplementary Table 1). The figure was created using modified PlotNeuralNet code [23].

**Supplementary Table 1:** Description of network blocks in tabular format. Parameters that were not specifically listed were set to the default values of TensorFlow version 2.2.

(a) List of parameters in every "Conv2D" layer.
| Parameter | Value |
| --- | --- |
| Kernel | 3x3 |
| Kernel initializer | He et al. |
| Padding | Same |
| Bias regularizer | L2 $10^{-6}$ |
| Kernel regularizer | L2 $10^{-6}$ |

(b) Layers in the "Convolutional block".
| Layer name | Operator | Input |
| --- | --- | --- |
| conv2d_1 | Conv2D | input |
| activation_1 | LReLU | conv2d_1 |
| conv2d_2 | Conv2D | activation_1 |
| activation_2 | LReLU | conv2d_2 |
| conv2d_3 | Conv2D | activation_2 |
| activation_3 | LReLU | conv2d_3 |

(c) Layers in the "Squeeze and excitation block".
| Layer name | Operator | Input |
| --- | --- | --- |
| gap2d | GlobalAveragePooling2D | input |
| dense_1 | Dense(units = $\max(n//8, 1)$) | gap2d |
| activation_1 | LReLU | dense_1 |
| dense_2 | Dense(units = n) | activation_1 |
| activation_2 | Sigmoid | dense_2 |
| multiply | Multiply | activation_2, input |

**Supplementary Figure 2:**
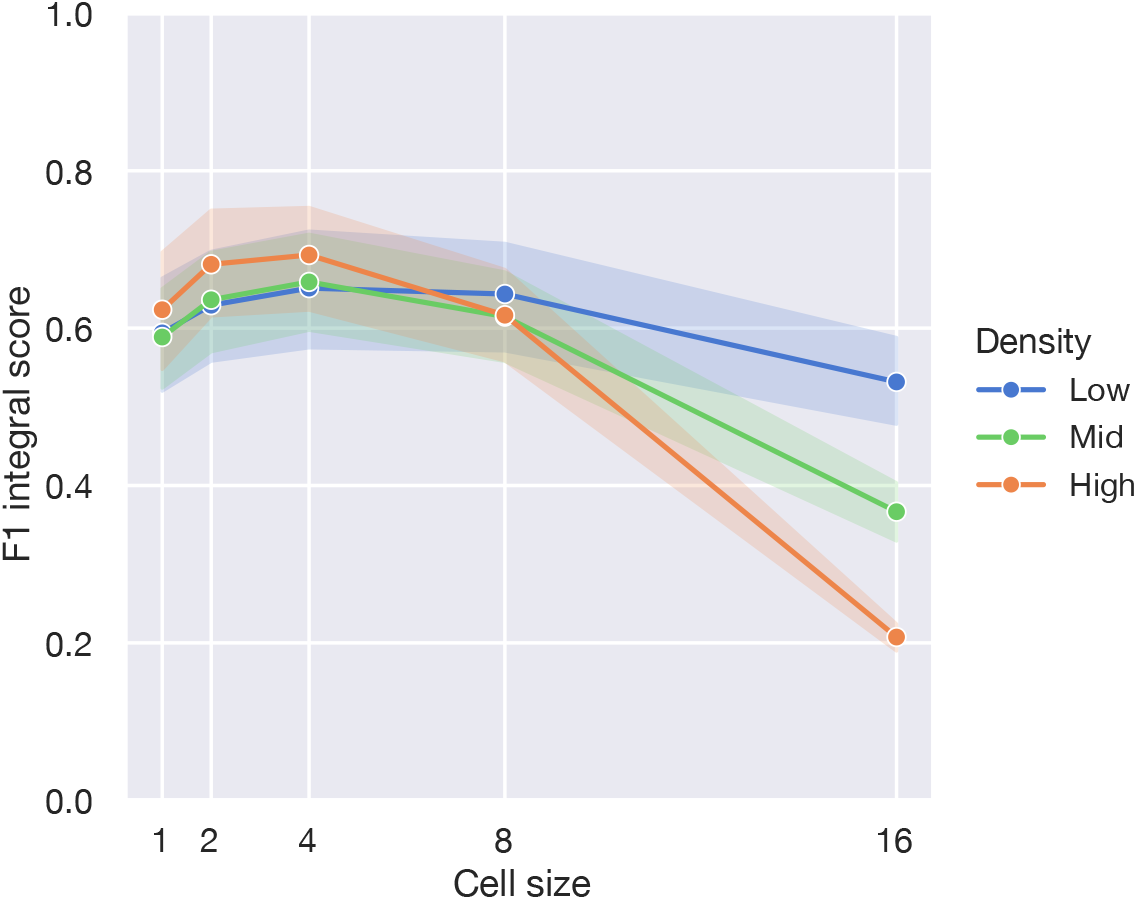
Relationship between cell size and F1 integral score at low, mid and high spot densities for the Receptor dataset.

**Supplementary Table 2:** Ablation table on the smFISH and Receptor datasets showing F1 integral scores (mean ± standard deviation) and percentage decrease (%-dec) upon removal of singular features. Features were sorted in descending order of mean percentage decrease. The “Replacement” column describes how each feature was replaced.

|  | Replacement | smFISH |  | Receptor |  | Mean |
| --- | --- | --- | --- | --- | --- | --- |
|  |  | Score | %-dec | Score | %-dec | %-dec |
| Full network |  | **0.905 ± 0.145** |  | **0.683 ± 0.309** |  |  |
| Dice Loss | BCE | 0.806 ± 0.196 | 10.990 | 0.639 ± 0.347 | 6.460 | 8.730 |
| RMSE Weight | $1/10$ | 0.856 ± 0.129 | 5.450 | 0.675 ± 0.304 | 1.160 | 3.310 |
| Squeeze Block | Remove | 0.893 ± 0.140 | 1.300 | 0.656 ± 0.317 | 3.930 | 2.620 |
| Filters $2^n$ | 4 | 0.885 ± 0.145 | 2.200 | 0.673 ± 0.301 | 1.450 | 1.830 |
| Spatial Dropout | Dropout | 0.900 ± 0.143 | 0.610 | 0.666 ± 0.309 | 2.430 | 1.520 |
| RMSE Weight | $1/2$ | 0.890 ± 0.135 | 1.670 | 0.678 ± 0.307 | 0.700 | 1.190 |
| Filters $2^n$ | 5 | 0.890 ± 0.144 | 1.610 | 0.680 ± 0.309 | 0.450 | 1.030 |
| Bottom Skip Connection | Remove | 0.895 ± 0.170 | 1.120 | 0.680 ± 0.308 | 0.370 | 0.750 |
| Batch Size | 1 | 0.900 ± 0.148 | 0.570 | 0.677 ± 0.309 | 0.850 | 0.710 |
| Filter Arrangement | Symmetrical | 0.897 ± 0.141 | 0.850 | 0.679 ± 0.309 | 0.560 | 0.710 |
| Convolutional Blocks | Residual | 0.903 ± 0.147 | 0.210 | 0.675 ± 0.311 | 1.080 | 0.650 |
| Optimizer | ADAM | 0.894 ± 0.154 | 1.190 | 0.682 ± 0.312 | 0.060 | 0.630 |
| L2 Regularization | Remove | 0.896 ± 0.165 | 1.020 | 0.682 ± 0.310 | 0.060 | 0.540 |
| Conv2D Activation | ReLu | 0.897 ± 0.153 | 0.900 | 0.683 ± 0.312 | -0.010 | 0.450 |
| Convolutional Blocks | Inception | 0.898 ± 0.137 | 0.740 | 0.683 ± 0.309 | 0.000 | 0.370 |
| Filters $2^n$ | 7 | 0.903 ± 0.146 | 0.200 | 0.680 ± 0.312 | 0.400 | 0.300 |
| Batch Size | 4 | 0.902 ± 0.150 | 0.330 | 0.682 ± 0.309 | 0.090 | 0.210 |

**Supplementary Figure 3:**
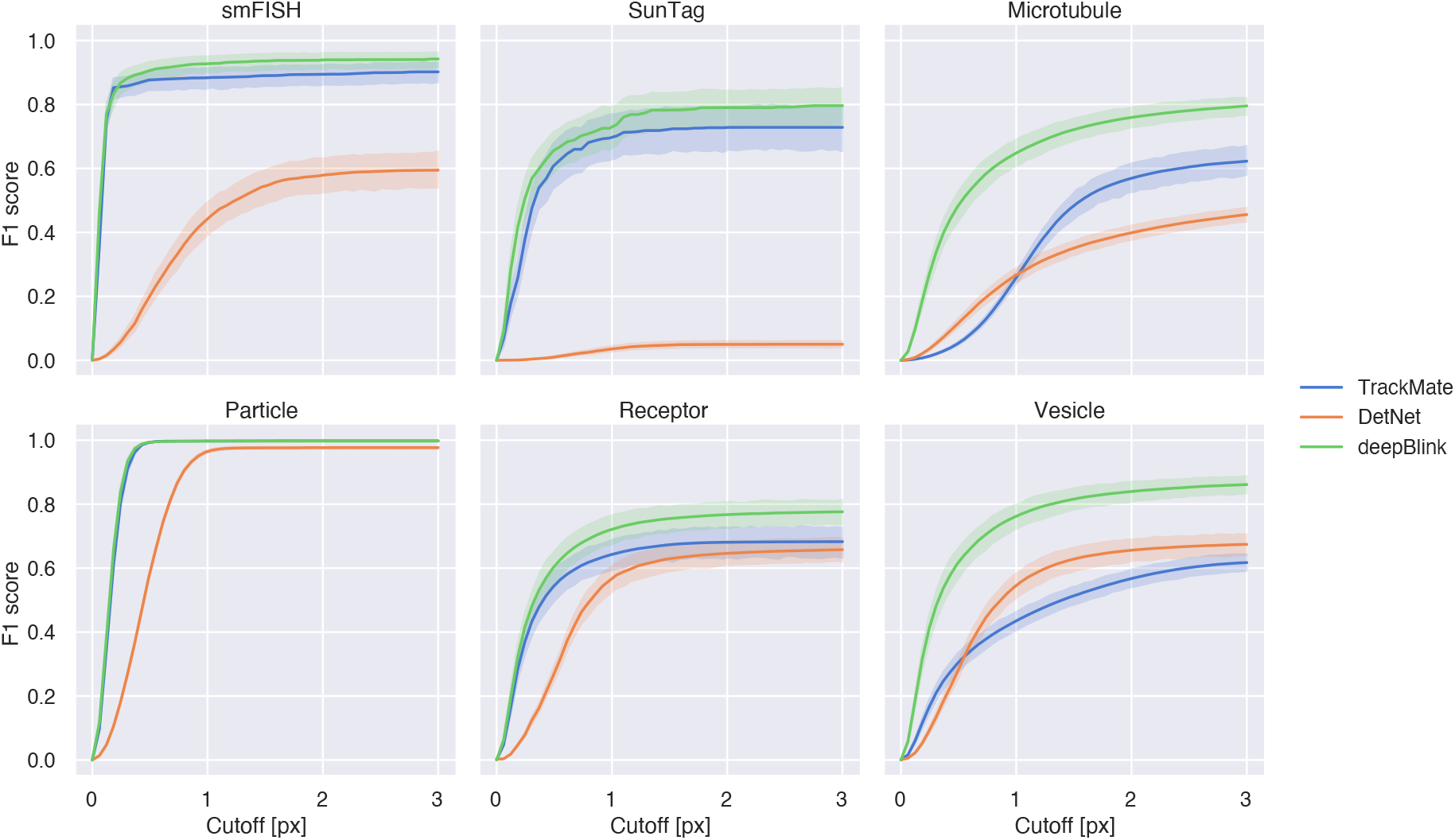
F1 score as a function of cutoffs on all datasets across and methods. Shaded areas correspond to the confidence interval of 95%.

**Supplementary Figure 4:**
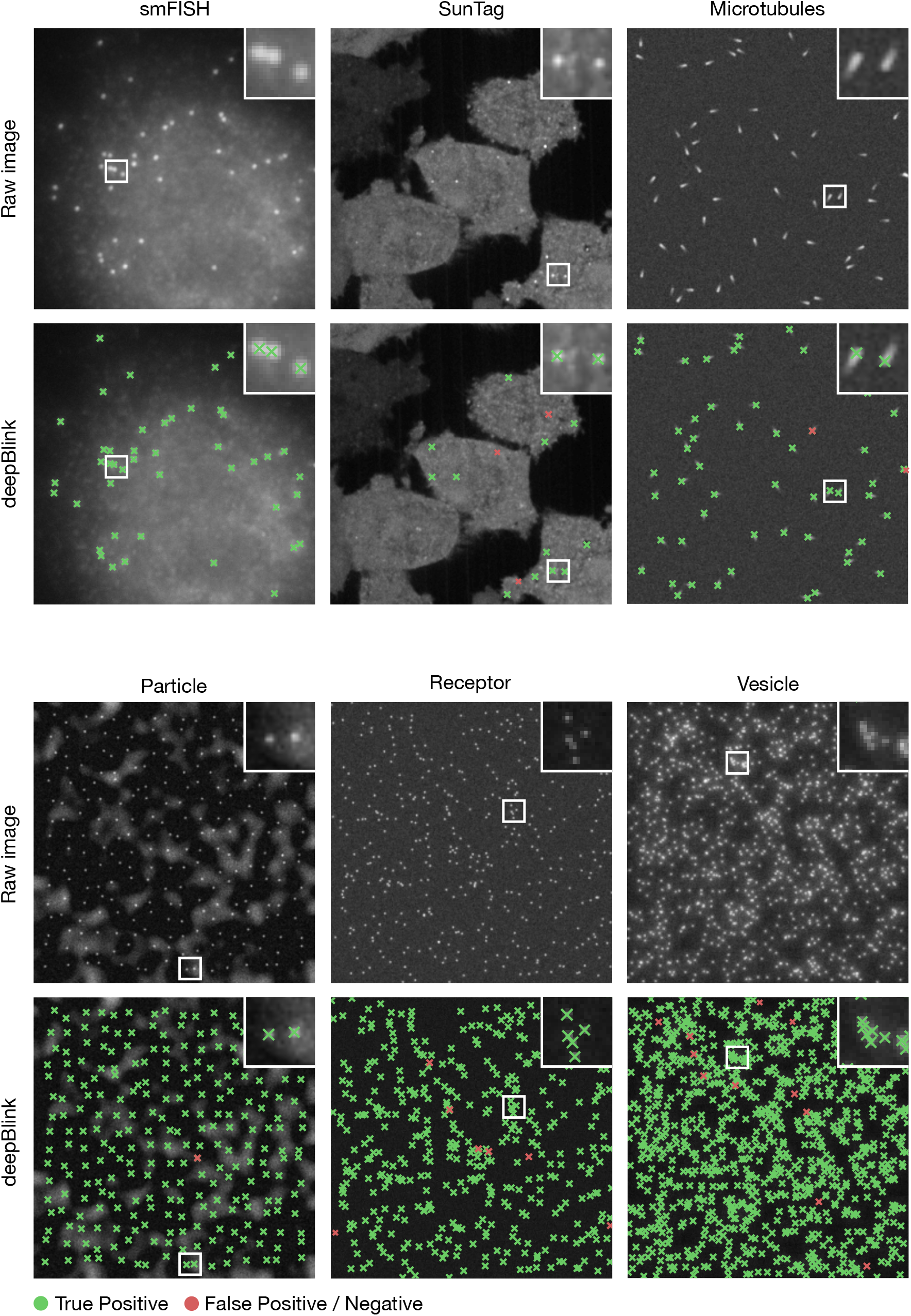
Representative images for all six datasets with their corresponding prediction.

**Supplementary Figure 5:**
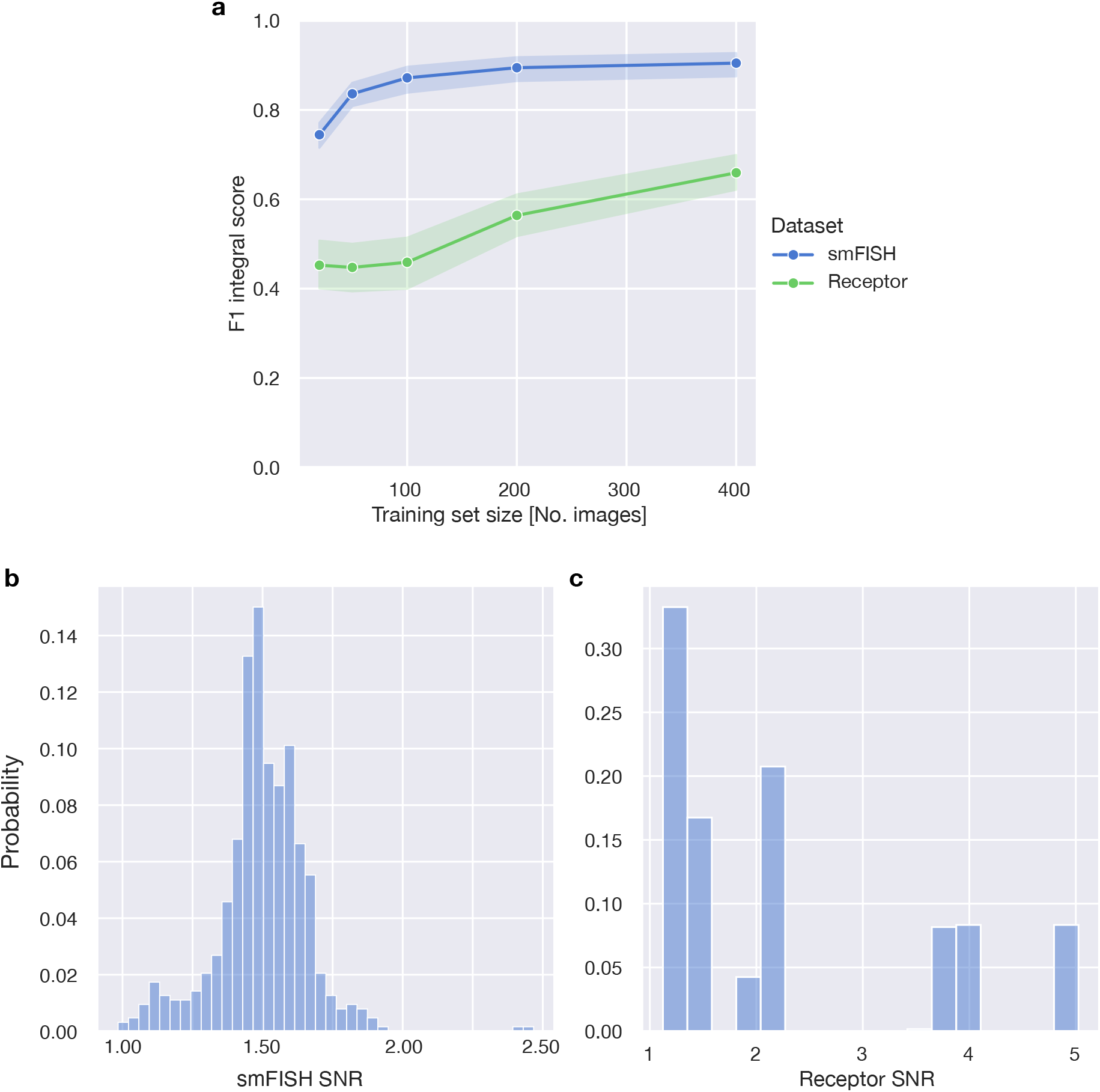
Effect of dataset size on model performance. **a** Relationship between training set size and F1 integral score on the same holdout test set. Shown for the smFISH and Receptor datasets. **b**, **c** show the SNR distribution across the entire dataset for smFISH and Receptor respectively.

**Supplementary Table 3:**
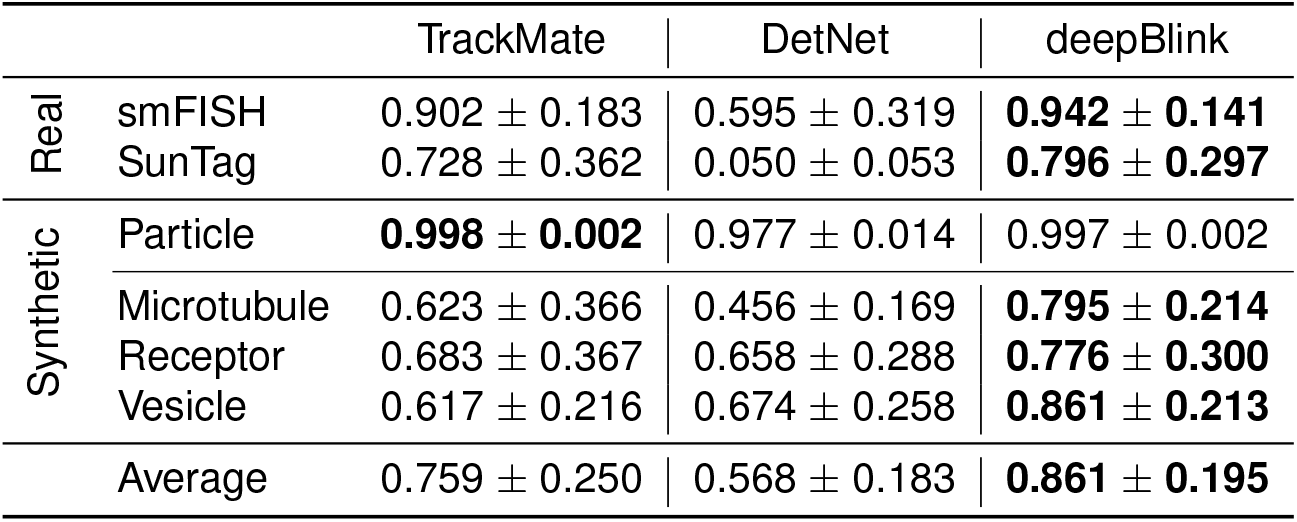
F1 scores at 3px cutoff of deepBlink and benchmarking methods (mean ± standard deviation). Shows values for three methods across six datasets. Higher value is better.

**Supplementary Table 4:** RMSE values of deepBlink and benchmarking methods (mean ± standard deviation). RMSE was only calculated on true positive spots. Shows values for three methods across four synthetic datasets, where the exact spot position is known. Lower value is better.

|  | TrackMate | DetNet | deepBlink + Gaussian | deepBlink |
| --- | --- | --- | --- | --- |
| Particle | 0.167 ± 0.020 | 0.438 ± 0.016 | 0.207 ± 0.021 | **0.157 ± 0.020** |
| Microtubule | 1.017 ± 0.221 | 0.887 ± 0.228 | 0.850 ± 0.223 | **0.588 ± 0.339** |
| Receptor | **0.428 ± 0.255** | 0.661 ± 0.203 | 0.517 ± 0.302 | 0.435 ± 0.294 |
| Vesicle | 0.658 ± 0.324 | 0.684 ± 0.223 | 0.644 ± 0.312 | **0.459 ± 0.311** |
| Average | 0.568 ± 0.205 | 0.667 ± 0.167 | 0.554 ± 0.215 | **0.410 ± 0.241** |

**Supplementary Figure 6:**
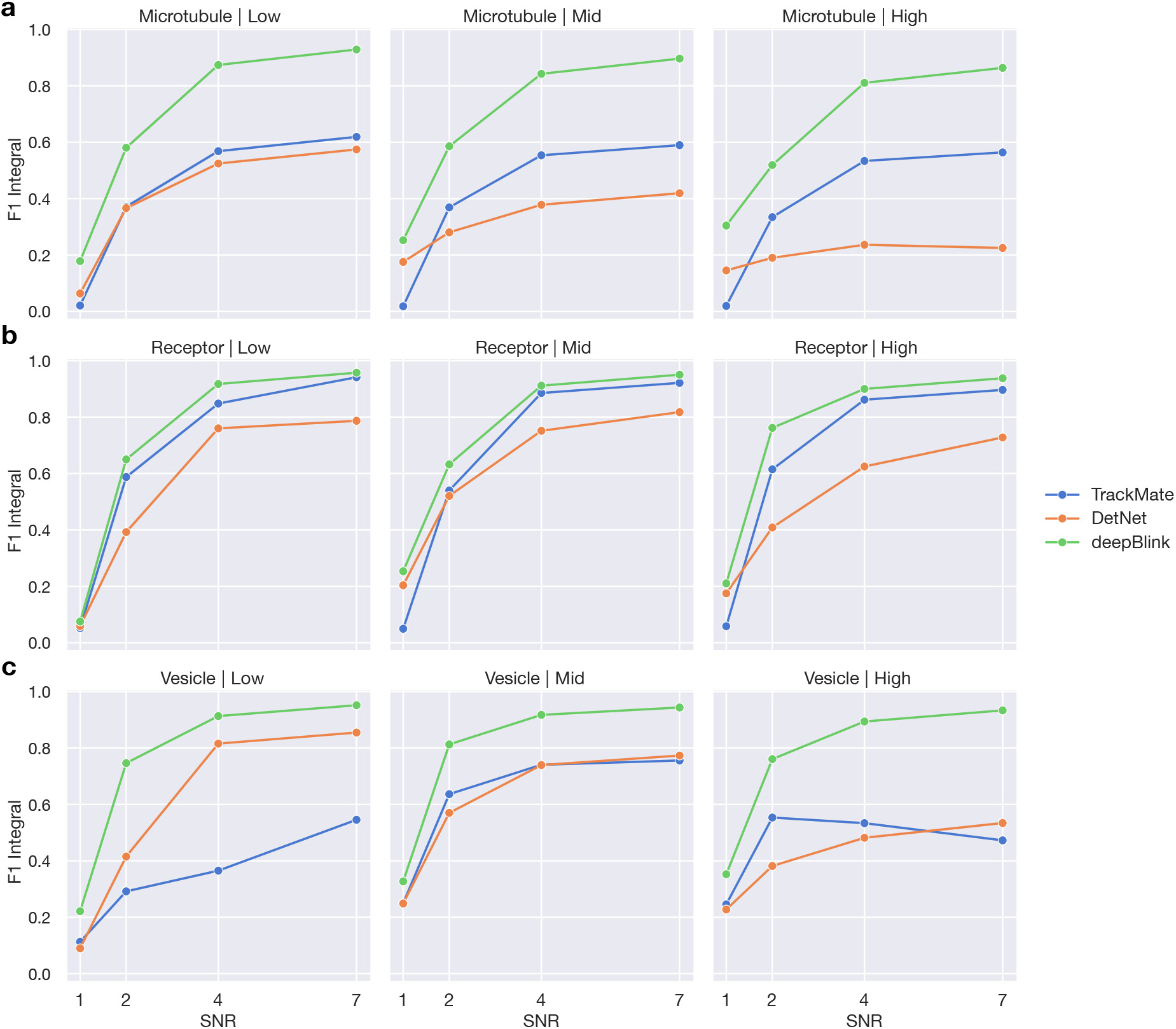
Mean F1 integral scores of all three methods at different spot densities and image SNRs for the **a** Microtubule, **b** Receptor, and **c** Vesicle datasets.

**Supplementary Figure 7:**
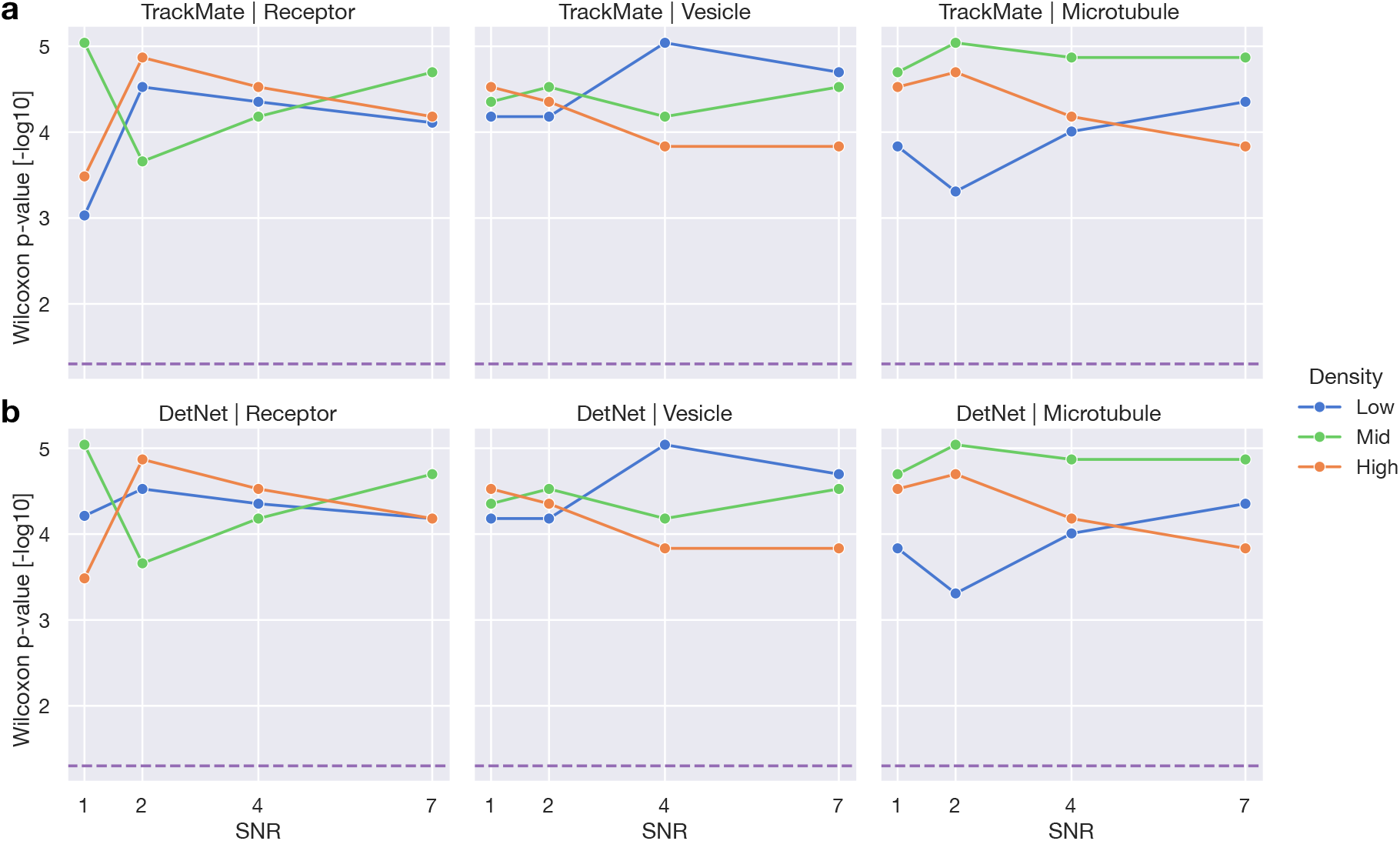
P-Values from one-sided Wilcoxon signed-rank tests measuring how significantly deepBlink performs better than **a** TrackMate and **b** DetNet at different spot densities and image SNRs. The purple dotted line indicates a significant p-value of 0.05.

**Supplementary Table 5:**
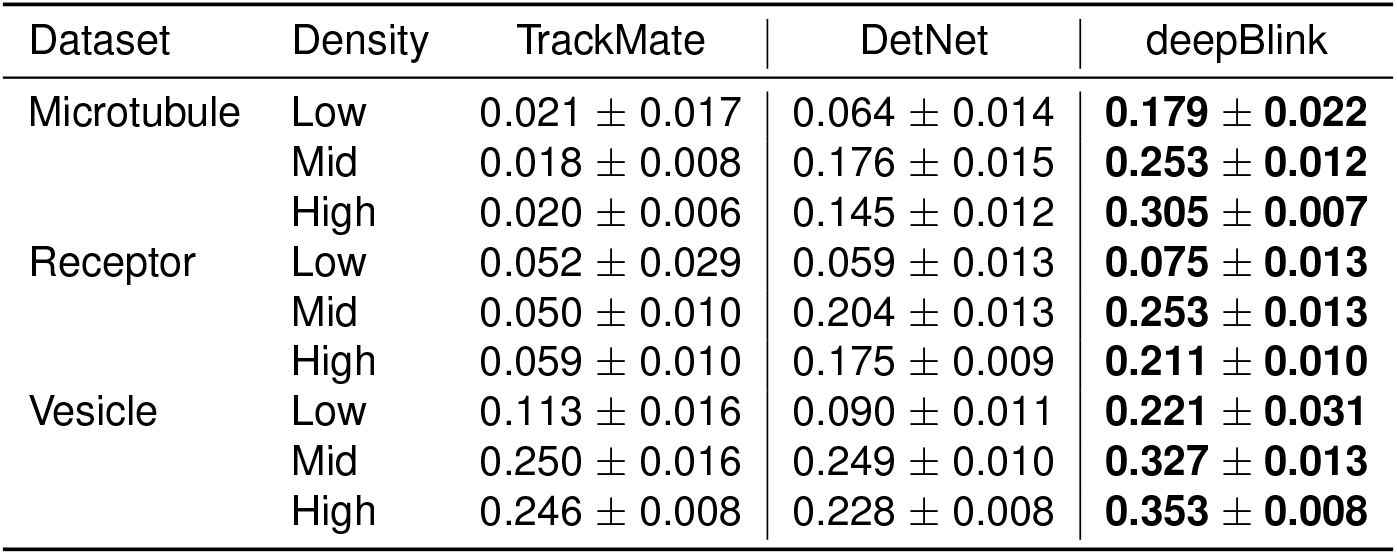
F1 integral score (mean ± standard deviation) for images of SNR equal to one. Shows values for three methods across three datasets.

**Supplementary Table 6:** The set of optimized parameters used for each benchmarking method and for each dataset.

(a) TrackMate
| Dataset | Diameter | Quantile |
| --- | --- | --- |
| smFISH | 4.0 | 0.871 |
| SunTag | 1.0 | 0.170 |
| Particle | 4.0 | 0.001 |
| Microtubule | 4.0 | 1.091 |
| Receptor | 3.0 | 0.795 |
| Vesicle | 3.0 | 1.342 |

(b) DetNet
| Dataset | Alpha |
| --- | --- |
| smFISH | 0.4737 |
| SunTag | 0.6842 |
| Particle | 0.5263 |
| Microtubule | 0.2105 |
| Receptor | 0.8946 |
| Vesicle | 0.5789 |

